# Astrocyte MCT1 expression does not contribute to the axonal degenerative phenotype observed with ubiquitous MCT1 depletion

**DOI:** 10.1101/2022.11.27.518094

**Authors:** Thomas Philips, Emily G. Thompson, Balaji G. Vijayakumar, Erica R. Kent, Sean J. Miller, Svetlana Vidensky, Mohamed Hassan Farah, Jeffrey D. Rothstein

## Abstract

We recently reported that loss of oligodendrocyte metabolic support through the lactate and pyruvate transporter Monocarboxylate Transporter 1 (MCT1) is well tolerated into adulthood. Only with advanced aging did we observe axonal degeneration and hypomyelination due to loss of MCT1 from oligodendroglia lineage cells. MCT1 is also expressed by other glial subtypes, such as astrocytes and endothelial cells where it has been suggested to be essential for learning and memory tasks. However, the importance of MCT1 in these cell types for long-term axonal metabolic support is still unknown. We therefore addressed whether conditional loss of MCT1 from either of these cell types would lead to widespread axonal degeneration with aging. Using a conditional null approach, similar to what was used for oligodendrocyte MCT1 depletion, we observed that conditional knockout of MCT1 from either astrocytes or endothelial cells did not cause neuronal injury. On the other hand, inducible ubiquitous depletion of MCT1 causes late-onset axonal degeneration, comparable with what was observed in our previous study using the oligodendrocyte lineage MCT1 null mice. In summary, we conclude that unlike oligodendrocyte MCT1, astrocyte MCT1 is not an essential driver of astrocyte mediated axonal energy homeostasis with aging.

## 1. Introduction

Glial cells in the central nervous system (CNS) express monocarboxylate transporters (MCTs) that allow for the passive transfer of energy metabolites like pyruvate and lactate across plasma membranes(Halestrap and Wilson, 2012). Various subtypes of MCTs are expressed in the CNS, with MCT1 and MCT4 most commonly found in glia while MCT2 is expressed by neurons(Pierre et al., 2000). It is now well established that neurons, especially those with long axons or complex dendritic arborizations, are highly dependent on the trophic energy mediators like glucose and lactate provided by glia(Suzuki et al., 2011, Tekkok et al., 2005, Lee et al., 2012, Funfschilling et al., 2012). These metabolic demands are met by lactate and pyruvate produced in astrocytes and oligodendrocytes during glycolysis, which can then be shuttled from glia towards neurons through their respective MCT subtypes. This provision of energy intermediates to neurons is integral to the so-called astrocyte-neuron and oligodendrocyte-neuron lactate shuttle where neuronal activity drives glial glucose metabolism and culminates in the shuttling of energy metabolites to neurons via MCTs(Pellerin and Magistretti, 1994, Saab et al., 2016, Philips and Rothstein, 2017). Upon transfer through neuronal MCTs, lactate and pyruvate are metabolized through oxidative phosphorylation to ensure the generation of sufficient levels of ATP, as neurons are unable to maintain energic demands solely through glycolysis (Almeida et al., 2001, Almeida et al., 2004). In addition, lactate has well-described alternative, non-metabolic functions that modulate neuronal excitability and plasticity upon transfer through MCTs or binding to lactate receptors (reviewed in (Magistretti and Allaman, 2018). The true directionality of lactate (from glia to neuron or the converse) is still under debate, as other studies suggest astrocytes import lactate through astrocyte MCT1 in complex with carbonic anhydrase 2 increases astrocyte glycogenolysis. The resulting glucose production in astrocytes drives glucose export through glucose transporters to neurons (Roosterman and Cottrell, 2020). Regardless of directionality, there is abundant evidence to suggest that the lactate transporter MCT1 is critical to providing neuronal metabolic support. Firstly, loss of MCT1 in ubiquitous heterozygous MCT1 null mice causes spatial memory deficits and late-onset axonal degeneration in the optic nerve, the latter attributed in part to the loss of MCT1 expression in oligodendrocytes (Lee et al., 2012, Tadi et al., 2015). In addition, multiple studies have demonstrated that rats or mice are unable to learn either an inhibitory avoidance or spatial memory task when hippocampal MCT1, MCT2 or MCT4 expression is downregulated pharmacologically, using MCT targeting antisense oligonucleotides (ASOs), or with Cre-loxP technology(Suzuki et al., 2011, Newman et al., 2011, Netzahualcoyotzi and Pellerin, 2020). Interestingly, the effects observed with ASOs targeting MCTs were attributed to MCT1/4 loss in astrocytes, but as this approach targets all cell types expressing MCT1 in the hippocampus, other cell types could have contributed to these effects as well. This is partially answered by a follow-up study in mice in which targeting astrocyte MCT4 expression led to spatial memory impairment (Netzahualcoyotzi and Pellerin, 2020). These studies are in line with those demonstrating an important role for glycogen-derived lactate in memory formation: targeting glycogenolysis with DAB in hippocampal astrocytes affects not only learning in rats but also learning-dependent synapse stabilization and changes in mRNA translation. These effects could then be rescued by the co-administration of DAB with lactate or pyruvate(Vezzoli et al., 2019, Descalzi et al., 2019, Suzuki et al., 2011). These studies suggest an important role for glycogen-derived lactate in learning-induced translation and synapse stabilization underlying long-term memory formation(Descalzi et al., 2019, Vezzoli et al., 2019). The exact mechanisms involved still need to be elucidated, especially given the multitude of non-metabolic contributions of lactate to neuronal excitability(Yang et al., 2014, Lauritzen et al., 2014)(reviewed in (Magistretti and Allaman, 2018)).

Lastly, MCT1 is strongly expressed by endothelial cells. Its role has been well studied in the context of tumor angiogenesis where targeting MCT1 affected lactate-induced HIF-1a activation and angiogenesis (Sonveaux et al., 2012). The importance of endothelial MCT1 in the CNS under homeostatic conditions is unknown. It has been suggested that endothelial MCT1 controls extracellular lactate levels during hippocampal neurogenesis, but whether CNS loss of endothelial MCT1 has any pathological consequences with aging remains to be explored (Wang et al., 2019). Interestingly, to our knowledge no studies in the past have addressed the importance of astrocyte and endothelial MCT1 with advanced aging. Therefore, to fully understand the importance for astrocyte MCT1 in fueling neuronal energy demands in the aging CNS, we developed conditional MCT1 null mice that were crossed with astrocyte-specific GFAP^Cre^ mice. To our surprise, we did not observe a degenerative phenotype when MCT1 was selectively depleted from astrocytes, even in mice aged to 2 years. Similar results were obtained when MCT1 was conditionally depleted from endothelial cells. These results suggest that compensatory astrocyte metabolic pathways are at play to ensure metabolic support to neurons is being maintained, or that MCT1 expression in other cell types can compensate for the loss of astrocytic MCT1. To elucidate the latter hypothesis, we utilized an approach to ablate MCT1 expression ubiquitously rather than in a cell-type specific manner. As MCT1 null mice die *in utero*, we used a conditional approach to delete MCT1 only when mice have reached adulthood. Interestingly, we observed that ubiquitous MCT1 depletion did cause late onset axonal injury as observed in mouse optic nerve, similar to the degeneration we recently reported upon conditional depletion of MCT1 from oligodendrocytes(Philips et al., 2021). In conclusion, we observe severe axonal degeneration with either oligodendroglial or ubiquitous inducible loss of MCT1. This data therefore suggests that MCT1 has an important role in axonal energy homeostasis with aging, but MCT1 expression in astrocytes and endothelial cells is dispensable for maintenance of axonal energy balance.

## 2. Materials and methods

### 2.1 Mice

B6.Cg-Tg(Gfap-cre)77.6Mvs/2J (Stock 012866, hereafter mGFAP^Cre^), B6.Cg-Tg(Tek-cre)1Ywa/J (Stock 008863, hereafter Tie2-Cre), B6.129X1-*Gt(ROSA)26Sor^tm1(EYFP)Cos^*/J (Stock 006148, hereafter RosaYFP) and Tg(CAG-cre/Esr1*)5Amc/J (Stock 004453, hereafter CAGGCreER) mice were purchased from Jackson Laboratories. *Mct1* transgenic mice in which exon2 of the mouse *Mct1* gene is inserted between two loxP sites (hereafter MCT1^lox^) were generated in our laboratory as described previously(Jha et al., 2020). Both male and female mice were used throughout our study and randomly allocated to each of the experimental groups at all ages. All mice were kept in 14 hours light/10hours dark cycle. All animal procedures were carried out in compliance with animal protocols approved by the Animal Use Committee at the Johns Hopkins University School of Medicine.

### 2.2 Crossing of transgenic mouse lines

The mGFAPCre mice were crossed with RosaYFP mice to generate double transgenic mGFAP^Cre^-RosaYFP animals, hemizygous for either transgene. The MCT1^lox^ conditional null mice were crossed with hemizygous mGFAP^Cre^ mice, hemizygous Tie2^Cre^ mice or hemizygous CAGG^CreER^ mice to obtain either double transgenic mGFAP^Cre^-MCT1^lox^ mice, double transgenic Tie2^Cre^-MCT1^lox^ mice or double transgenic CAGG^CreER^-MCT1^lox^ mice, hemizygous for Cre and homozygous for the MCT1 conditional allele. All mice used in this study were hemizygous for their respective Cre and homozygous for the MCT1^lox^ allele.

### 2.3 Animal behavior and treatment procedures

All mouse behavior experiments were performed at the Behavioral Core of the Johns Hopkins School of Medicine.

#### Accelerated Rotarod

Motor learning and coordination were examined by the rotarod test (Rotamex-5 System with a spindle dimensions of 3.0 cm x 9.5 cm; Columbus Instruments). Testing was conducted over a 3 day period, where each mouse was given three trials per day and latency to fall off the rotarod was measured as it accelerated from 4 to 99 rpm over a 5 min. period. Before the start of testing on the first day, each mouse was given a habituation trial by being placed on the rotarod, which was rotating at a constant speed of 4 rpm for 10 min. The mean latencies were calculated for each animal.

#### Open field testing

Spontaneous locomotion was assessed over a 30 min. period using activity chambers (16” (W) x 16” (D) x 15” (H)) with Photobeam Activity System (PAS) open field system equipped with 16 x 16 infrared beams (San Diego Instruments, San Diego, CA, USA) during the dark period from 10:00am to 06:00pm (reversed light cycle). Total locomotor activity was automatically measured as the number of beams breaks. The center zone was defined as the surface area covered by the central 14 x 14 beams of the box (14 × 14 inches) with the remaining peripheral area being designated as the peripheral zone. The amount of time mice spend along the walls was used as a measure of anxiety.

#### Y-maze

Spatial working memory assessed by spontaneous alternation and spatial recognition memory were assessed in the y-maze test as previously described(Shevelkin et al., 2017).

#### Passive avoidance test

Passive avoidance was assessed using GEMINI Active and Passive Avoidance System (Stoelting Co., Wood Dale, IL). Briefly, a mouse was placed in the light compartment of the shuttle box for 30 sec. to acclimate. Following the habituation, the door between two compartments was opened, and the latency to enter the dark compartment was measured. Entering the dark compartment led to the door closing. Three seconds after entering the dark compartment, the mouse was exposed to an inescapable foot shock delivered at the intensity of 0.5mA. Twenty-four hours following the training session, the mouse was returned to the light compartment, the door between the compartments was opened and the latency to enter the dark compartment was measured.

#### Elevated Maze Plus

Anxiety was evaluated in the plus-maze (San Diego Instruments Inc., San Diego, CA, USA) as previously described(Pletnikov et al., 2008).

#### Trace fear trace conditioning test

Trace fear conditioning (TFC) was assessed as previously described(Jouroukhin et al., 2018, Terrillion et al., 2017). Briefly, TFC is a 3 day test consisting of the habituation day, training day, and test day. On day 1, a mouse was habituated to the shock box (Coulbourn, Holliston, MA) for 10 min. On Day 2, the mouse was placed in the shock box, and a 20 sec. 90dB tone was delivered. Twenty seconds following the termination of the tone, a scrambled 2 sec. 0.5 mA shock was delivered. This tone-shock pairing was repeated three times. On Day 3, the mice were placed in the shock box for 3 min and the freezing behavior was assessed as a measure of contextual memory. Following the context test, the cued test was performed. The mice were placed in a different testing box, and the tone presented during the training session was presented three times. The freezing behavior in response to the tones was assessed as a measure of cue-dependent fear memory. Freezing was automatically detecting using the Freeze Scan software (Clever Sys Inc.).

### 2.4 Tamoxifen injections

Tamoxifen (MilliporeSigma) was dissolved in ethanol, vortexed for 10 sec. and dissolved in sunflower seed oil (ratio ethanol-oil 1:10). CAGG^CreER^ mice crossed with MCT1^lox^ mice were injected intraperitoneally with 4 doses of tamoxifen (200mg/kg) for 4 consecutive days at the age of 30 days.

### 2.5 Immunohistochemistry

Mice were deeply anesthetized by I.P. injection of an anaesthetic cocktail (hereafter KXA) including: acepromazine maleate (2.5mg/kg; MWI Veterinary Supply), ketamine (100mg/kg; MWI Veterinary Supply), and xylazine (10mg/kg; MWI Veterinary Supply). Mice were transcardially perfused with ice-cold PBS followed by ice-cold PBS containing 4% paraformaldehyde (PFA). Spinal cord, cerebellum, optic nerve and forebrains were dissected and post-fixed in 4% PFA in PBS for 4 hours. Samples were stored overnight in 30% sucrose solution in PBS at 4°C and subsequently embedded in OCT compound (Sakura Finetek). 20 µm spinal cord and 40 µm coronal brain and cerebellar floating sections were cut on a cryostat (Thermo Scientific), washed in PBS and blocked for 1 hour at room temperature in PBS solution containing 0.1% Triton X-100 (PBST) and 10% normal goat serum (Vector). Sections were then incubated overnight at 4°C with primary antibodies diluted in PBST: chicken ant-MCT1 (our lab), rabbit anti-Iba1 (WAKO), chicken anti-GFAP (MilliporeSigma), goat anti-Chat (MilliporeSigma), rabbit anti-GFP (Rockland), rabbit anti-GLT-1 (our lab) and mouse anti-NeuN (MilliporeSigma). Sections were washed in PBST and incubated with fluorescent labeled secondary antibodies (Thermo Scientific) for 1 hour at room temperature. Sections were subsequently washed in PBST and mounted on glass slides in Prolong Gold antifade reagent containing DAPI (Thermo Scientific). Sections were imaged on a Zeiss Axioimager Z1 equipped with Apotome 2 or on a Zeiss 800 confocal microscope (Zeiss).

### 2.6 Myelin preparation

Myelin was isolated and purified as reported earlier (Erwig et al., 2019). In brief, whole brain and spinal cord were dissected and homogenized in 0.32M sucrose using an ultra-turrax homogenizer. The homogenate was carefully layered over a 0.85M sucrose solution and centrifuged at 75000g for 30 min. at 4°C in a Beckman ultracentrifuge (Beckman). After centrifugation, the myelin containing interface was collected, washed with deionized water, and centrifuged at 75000g for 15 min at 4°C. The pelleted fractions then underwent two rounds of osmotic shock by incubation in deionized water for 10 min. on ice followed by ultracentrifugation at 12000g for 15 min. at 4°C. Pellets were then dissolved in 0.32M sucrose, layered over 0.85M sucrose and the myelin containing interface was collected after centrifugation at 75000g for 30 min. at 4°C. After another wash in deionized water, myelin was pelleted by centrifugation at 75000g for 15 min at 4°C. Myelin pellets were dissolved in RIPA buffer (Thermo Scientific) containing protease and phosphatase inhibitors and stored at −80°C.

### 2.7 Western Blot

The protein concentration of each sample was measured using the Lowry method and a SpectraMax M3 plate reader (Molecular Devices) was used to measure the colorimetric reaction. 10-40µg of total protein was loaded on mini Protean TGX Precast gels (Biorad). After electrophoresis, proteins were blotted on a PVDF membrane (Biorad) and blocked in 5% Blotting Grade Blotter (Biorad) diluted in Tris buffered saline (TBS) containing 0.1% Tween 20 (TBST). Blots were then incubated with either of the following antibodies: mouse anti-PLP/DM20 (MillipreSigma), chicken anti-human MCT1 antibody (our lab), mouse anti-GAPDH (Thermo Scientific). Membranes were incubated overnight with primary antibodies at 4°C. Membranes were then washed with TBST and incubated with HRP labeled secondary antibodies for 1 hour at room temperature. After two washes in TBST and one wash in TBS, membranes were exposed to ECL reagent for 2 min. (Thermo Scientific) and chemiluminescent signal was detected with the LAS Imager 4000 (GE Healthcare).

### 2.8 Lectin injections

Tie2^Cre^MCT1^lox^ and MCT1^lox^ littermate control mice were deeply anesthetized with KXA as written above. Right ventricles were injected with 100ul of 1ug/ul Lycopersicon Esculentum (Tomato) Lectin (LEL, TL), DyLight 488 (Thermo Scientific) solution. Five minutes after injection, brains were dissected and flash frozen in isopentane chilled with liquid nitrogen. 20um coronal brain sections were cut on a cryostat and green fluorescence was analyzed on a Zeiss Apotome Imager Z1.

### 2.9 Spine density analysis

Whole brains from P360 mGFAP^Cre^-MCT1^lox^ and MCT1^lox^ littermate controls were stained with a Golgi-Cox stain kit (FD Rapid GolgiStain, FD NeuroTechnologies, Inc. PK401A) according to the manufacturer’s instructions. Golgi-Cox stained 100μm coronal brain slices were then cut on a cryostat and imaged on a Zeiss Apotome Imager Z1 equipped with a 100X objective. Blinded spine density analysis was performed using Neurolucida 360 (MBF Bioscience) software. Spine density analysis was completed for 3-5 mice of each genotype in both the cortex (7-10 spines per animal) and the hippocampus (4-9 spines per animal).

### 2.10 Reverse transcriptase-q-PCR

Mice were deeply anesthetized with KXA as written above and transcardially perfused with ice-cold PBS. Tissue was immediately dissected and submerged in Tri Reagent (Zymo Research). Alternatively, oligodendrocyte lineage cells were sorted from whole tissue as described above. RNA was isolated using either the Direct-zol RNA MicroPrep kit (Zymo Research) (for cells) or by conventional chloroform-isopropanol-ethanol extraction methods (for tissue). For the latter, tissue in Tri Reagent was diluted in chloroform in a 4:1 Tri Reagent:chloroform ratio. After centrifugation at 12000rpm, the clear upper phase is collected, and ice-cold isopropanol is added. After 12 minutes centrifugation at 12000rpm, 75% ethanol is added to the pellet. After subsequent centrifugation (12000rpm, 10 minutes), the RNA pellet is dried and dissolved in purified water. RNA concentration was measured by Nanodrop (Thermo Scientific). RNA then underwent reverse transcription into cDNA using the High Capacity cDNA reverse transcription kit (Thermo Scientific). To detect relative gene expression, inventoried assay Taqman probes targeting *Mct1, Mct2, Mct4, Cx43, Cx30, Cx47, Glut1*, *Glut3* and *Gapdh* (endogenous control) were used.

### 2.11 Electron microscopy

Mice were deeply anesthetized with KXA as written above and transcardially perfused with 0.1M Sorensen’s phosphate buffer and subsequently perfused with 4% paraformaldehyde/2.5% glutaraldehyde in 0.1M phosphate buffer. Optic nerves were dissected and post-fixed with 2% osmium for 2 hours. The tissue was then dehydrated in graded ethanol and embedded in Embed 812 resin (EMS). Thin sections (70-80nm) were cut on a Reichert Jung Ultracut E microtome and placed on formvar coated 100 mesh copper grids. The sections were stained with uranyl acetate followed by lead citrate. All imaging was performed on a Zeiss Libra 120 with a Veleta (Olympus) camera at an accelerating voltage of 120Kv. For light microscopy, 0.5µm sections were cut and stained with 1% toluidine blue (EMS).

### 2.12 Primary astrocyte cultures

Primary astrocyte cultures were prepared from P7 mouse cortices from mGFAP^Cre^-MCT1^lox^ and MCT1^lox^ mice. Both male and female pups were allocated to each experimental group. Cortices were dissociated using Miltenyi’s adult mouse brain dissociation kit according to the manufacturer’s protocol. In short, meninges were removed, and cortices underwent papain digestion for 30 min. using the adult brain dissociator (Miltenyi). After passage of cells through a 70µm cell strainer, myelin was removed with ‘myelin debris removal solution’ and single cells were pelleted. Red blood cells were removed using ‘Red Blood cell Removal Solution’ and cells were incubated with anti-Acsa1 magnetic microbeads for 15 minutes on ice. Acsa1+ astrocytes were then magnetically sorted and collected in astrocyte cell medium (DMEM-F12 with Hepes and Glutamax, 10% FBS and Pen/Strep). Cells were plated at T75 flasks coated with Matrigel and maintained in the cell incubator at 37.5°C and 5% CO_2_ until grown to a confluent monolayer for 10-14 days. Media changes was performed 24 hours after plating and then every fourth day.

### 2.13 Lactate and pyruvate transport assay

Lactate and pyruvate transport assays were performed on astrocyte primary cell cultures. After allowing isolated astrocytes to grow to confluency in T75 flasks for 10-14 days, cells were passaged to 12-well plates at 200K cells per well before lactate/pyruvate transport in astrocyte cultures was measured with the lactate/pyruvate transport assay. For measuring lactate/pyruvate uptake, cells were washed twice in HEBSS buffer (25mM Hepes, 150mM NaCl, 3mM KCl, 1.7mM KH_2_PO_4_, 0.6mM MgCl_2_ and 1mM CaCl_2_*2H_2_O). Cells were then incubated in HEBSS for 10 min on ice with 1uM of AR-C1554858 (Tocris) in HEBSS buffer or untreated. After incubation cells were exposed to 2X Hot Lactate/Pyruvate solution (0.5µCi/mL of ^14^C-L-lactate/^14^C Pyruvate, PerkinElmer) for 1 min. on ice. Cells were washed twice with HEBSS buffer containing 2mM 4-CIN and 0.1N NaOH was added to the wells. Plates were swirled to detach the cells from the bottom of the plate and cells were lysed by pipetting cells up and down 30 times. 300µl of the supernatant was then transferred to a scintillation vial and 3 ml of scintillation fluid was added to each sample. Samples were placed on a shaker overnight at room temperature and radioactivity of the samples was measured with a scintillation counting machine. The remaining supernatants obtained after cell lysis were used to measure protein concentration with a Bradford assay, where the colorimetric reaction was measured on a SpectraMax M3 plate reader (Molecular Devices).

### 2.14 Image and data analysis

Images were analyzed using Image-J bundled with Java 1.8.0_112 and Zen 2.3 Lite software. For immunofluorescence, at least 2-15 sections per mouse were analyzed and quantified using Zen 2.3 Lite or Image-J. Cells were quantified within a defined area as # cells/µm^2^. For electron microscopy images g-ratio’s and axonal degenerative phenotypes were measured on >500 axons per sample from 3-10 images taken at 10000x magnification. Images were taken from a minimum of 5 randomly chosen areas of an optic nerve cross section. Inner and outer area of axons were measured with Image J from which the outer and inner axonal diameter was calculated respectively. Axons were scored as degenerated if they were swollen with accumulation of intracellular material (vesicles, swollen mitochondria), a dark cytoplasm, or if axons had disappeared altogether with an accumulation of redundant myelin loops. Mitochondrial area was calculated with Image J bundled with Java 1.8.0_112.

### 2.15 Statistics

All data is represented as mean +/-SEM as indicated in the figure legends. All statistical details of the experiments can be found in either the Figure legends or the Supplementary Figure legends including the value of n, the p-value and statistical test being used. The p-value was considered significant when p<0.05. No subjects were excluded from the datasets in any of the experiments. All data met assumptions for the statistical approach, e.g. equal variance among groups. All statistical analysis was performed using GraphPad Prism7.

## 3. Results

### 3.1 Astrocyte MCT1 protein and transporter activity is reduced in astrocyte MCT1 conditional null mice

To knock out MCT1 from astrocytes specifically, we crossed B6.Cg-Tg(Gfap-cre)77.6Mvs/2J mice (GFAP^Cre^ mice) with the MCT1 conditional null mice (hereafter named MCT1^lox^) to obtain double transgenic GFAP^Cre^-MCT1^lox^ mice, hemizygous for Cre and homozygous for the MCT1^lox^ allele. Homozygote MCT1^lox^ littermates were used as controls. We first validated MCT1 knockout in the cortex and spinal cord of P30 GFAP^Cre^-MCT1^lox^ mice as compared with age-matched MCT1^lox^ controls. Quantitative PCR revealed a 24% reduction in MCT1 mRNA in the spinal cord whereas the expression of other MCTs like MCT2 and MCT4 was unaffected (Supplementary Figure 1B). Western Blot revealed a 67% reduction in MCT1 protein expression in spinal cord and a 33% reduction in the cortex (Figure 1A). In addition, we performed quantitative PCR to determine whether changes in the expression of glucose transporters or connexin hemichannels could metabolically compensate for loss of astrocyte MCT1 but found no significant differences in their expression (Supplementary Figure 1C). We performed MCT1 immunostaining of different CNS areas to confirm the loss of MCT1 expression in astrocytes (Figure 1B and Supplementary Figure 1A). In control tissue, MCT1 expression correlated strongly with GLT-1 expression in fine astrocyte processes. As expected, MCT1 expression had disappeared from the GLT-1 expressing astrocyte processes in GFAP^Cre^-MCT1^lox^ mice at both P30 and P360, the oldest age analyzed (Supplementary Figure 1A and Figure 1B). To determine if oligodendroglia MCT1 expression was affected by loss of astrocyte MCT1, we compared MCT1 expression in the myelin from GFAP^Cre^-MCT1^lox^ and age-matched control mice and found no significant difference (Supplementary Figure 1D). Lastly, to confirm that we had obtained functional loss of astrocyte MCT1 transporter activity in the GFAP^Cre^-MCT1^lox^ conditional null mice, we performed radioactive ^14^C lactate and pyruvate uptake assays. Astrocytes were cultured from P3 MCT1^lox^ or GFAPCre-MCT^1lox^ transgenic pups and either ^14^C lactate or ^14^C pyruvate were added to the culture medium to measure substrate uptake. We observed a 61% reduction in ^14^C lactate uptake in the astrocyte cultures derived from the GFAP^Cre^-MCT1^lox^ mice as compared to the MCT1^lox^ controls, which was similar to what was observed with the MCT1 inhibitor AR-C155858 (Figure 1C). When measuring ^14^C pyruvate uptake, we observed a 48% reduction in the mGFAP^Cre^-MCT1^lox^ mice as compared to the MCT1^lox^ controls (Figure 1D). When using the MCT1 inhibitor AR-C155858 on the MCT1^lox^ cultures, we observed an 80% decrease in uptake of radioactive pyruvate (Figure 1D). AR-C155858 had no effect on ^14^C pyruvate uptake measured in the mGFAP^Cre^-MCT1^lox^ cultures (Figure 1D). These data therefore validate the use of the mGFAP^Cre^-MCT1^lox^ mice for the study of behavioral and pathological changes in response to loss of astrocyte MCT1 expression.

**Figure 1:**
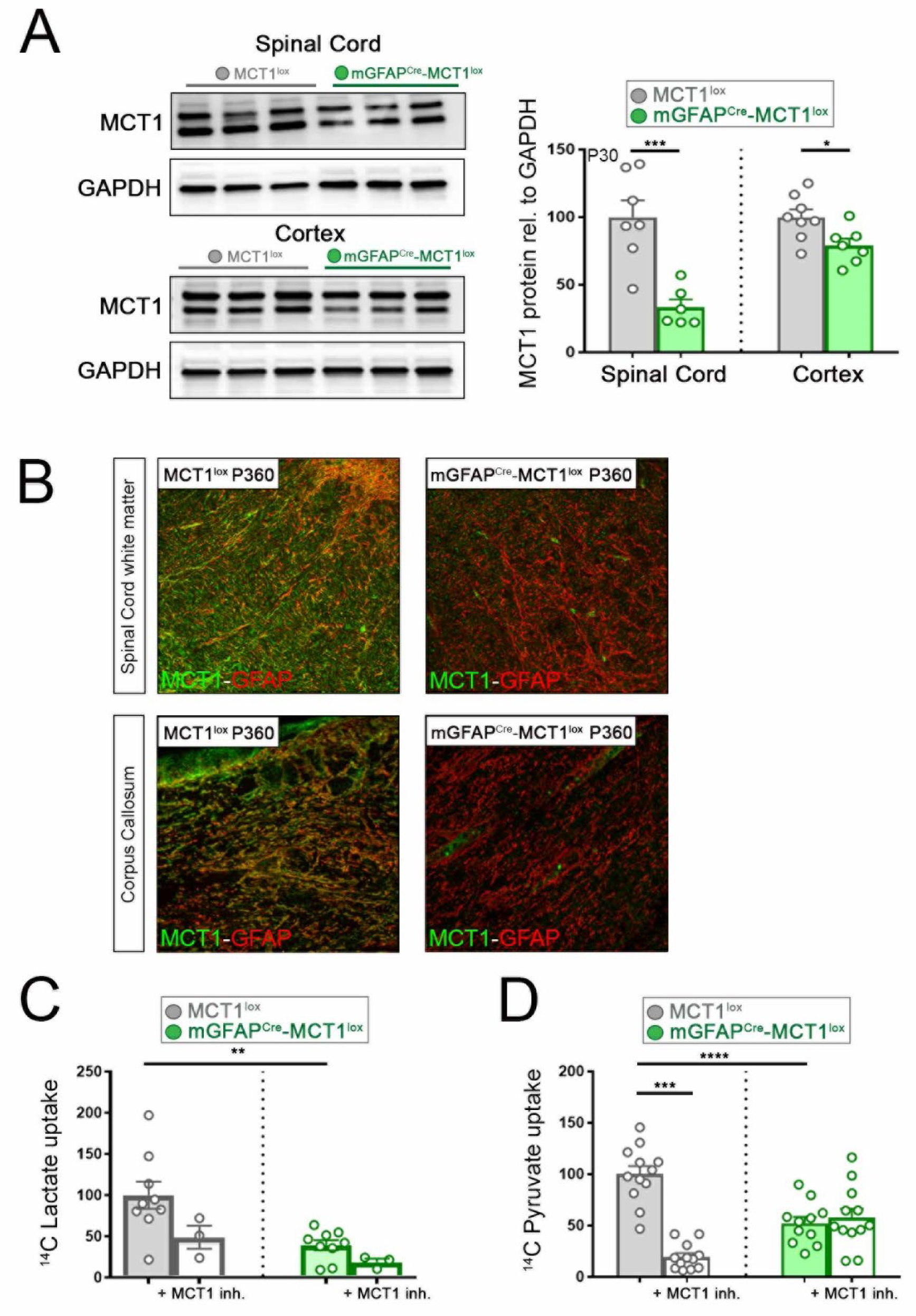
Loss of astrocyte MCT1 expression in astrocyte MCT1 conditional null mice. (A) MCT1 protein expression in P30 mGFAP^Cre^-MCT1^lox^ and MCT1^lox^ controls is reduced by 67% in the spinal cord (n=7-8, ***p<0.001, Student’s t-test) and 33% in cortex (n=7-8, and *p<0.05, Student’s t-test). Data is represented as mean ± SEM. (B) Immunostaining of MCT1 in the spinal cord (top) and corpus callosum (bottom) of P360 mGFAP^Cre^-MCT1^lox^ and MCT1^lox^ mice. Scale bar is 100µm. Images are a representative a minimum of n=3. (C) Radioactive lactate transport was reduced by 61% in mGFAP^Cre^-MCT1^lox^ primary astrocyte cultures as compared to MCT1^lox^ cultures (n=3-9, **p<0.01, one-way ANOVA). Data is represented as mean ± SEM. (D) Radioactive pyruvate transport was reduced by 48% in mGFAP^Cre^-MCT1^lox^ primary astrocyte cultures as compared to MCT1^lox^ cultures (n=12, ****p<0.01, one-way ANOVA). Data is represented as mean ± SEM.

### 3.2 Loss of astrocyte MCT1 does not lead to behavioral changes in mGFAP^Cre^-MCT1^lox^ mice

We first compared general metabolism between P60 GFAP^Cre^-MCT1^lox^ and MCT1^lox^ control mice. We analyzed oxygen consumption, CO2 production, respiratory exchange ratio and chow/water intake in the GFAP^Cre^-MCT1^lox^ mice as compared to MCT1^lox^ controls using the Comprehensive Lab Animal Monitoring System (CLAMS). We did not observe any significant differences between the two groups analyzed (data not shown). Additionally, we found no differences in the percentage of fat and lean mass or changes in total body weight between GFAP^Cre^-MCT1^lox^ mice and MCT1^lox^ controls (Supplementary Figure 2A-B). We then explored whether loss of astrocyte MCT1 resulted in motor function deficits or changes in exploratory behavior. We performed accelerated rotarod, open field test and Catwalk analysis on P60 GFAP^Cre^-MCT1^lox^ mice using age-matched MCT1^lox^ mice as controls. We did not observe significant differences in rotarod performance (Figure 2A) or changes in gait and locomotion upon Catwalk analysis (data not shown). There were also no changes in open field behavior, rearing activity or anxiety in GFAP^Cre^-MCT1^lox^ mice as compared to MCT1^lox^ control mice (Figure 2B-D). Females and males performed equally well for both genotypes analyzed. We then analyzed working memory with Y-maze (Figure 2E and Supplementary Figure 2C)and learning and memory with passive avoidance test (Figure 2F) but found no significant differences between the GFAP^Cre^-MCT1^lox^ and MCT1^lox^ control mice. We did observe a modest context-dependent memory deficit in the fear conditioning test suggesting that these mice have a decreased sensitivity to a strong anxiety stimulus (Figure 2G-H and Supplementary Figure 2D-E). As it has been reported that inhibition of astrocyte glycogenolysis affects hippocampal synaptic density(Vezzoli et al., 2019), an effect attributed to a reduction in astrocyte lactate production, we performed Golgi-Cox staining on coronal brain sections isolated from mGFAP^Cre^-MCT1^lox^ and MCT1^lox^ controls and quantified synaptic density in the cortex and the hippocampus. As shown in Figure 2I, we did not observe changes in synaptic density between mGFAP^Cre^-MCT1^lox^ and MCT1^lox^ controls, suggesting that MCT1 is dispensable for maintenance of synaptic density in animals without astrocytic MCT1 expression. In conclusion, apart from changes related to recognition memory, overall behavior in the GFAP^Cre-^MCT1^lox^ mice is unaffected when compared to age-matched MCT1^lox^ controls.

**Figure 2:**
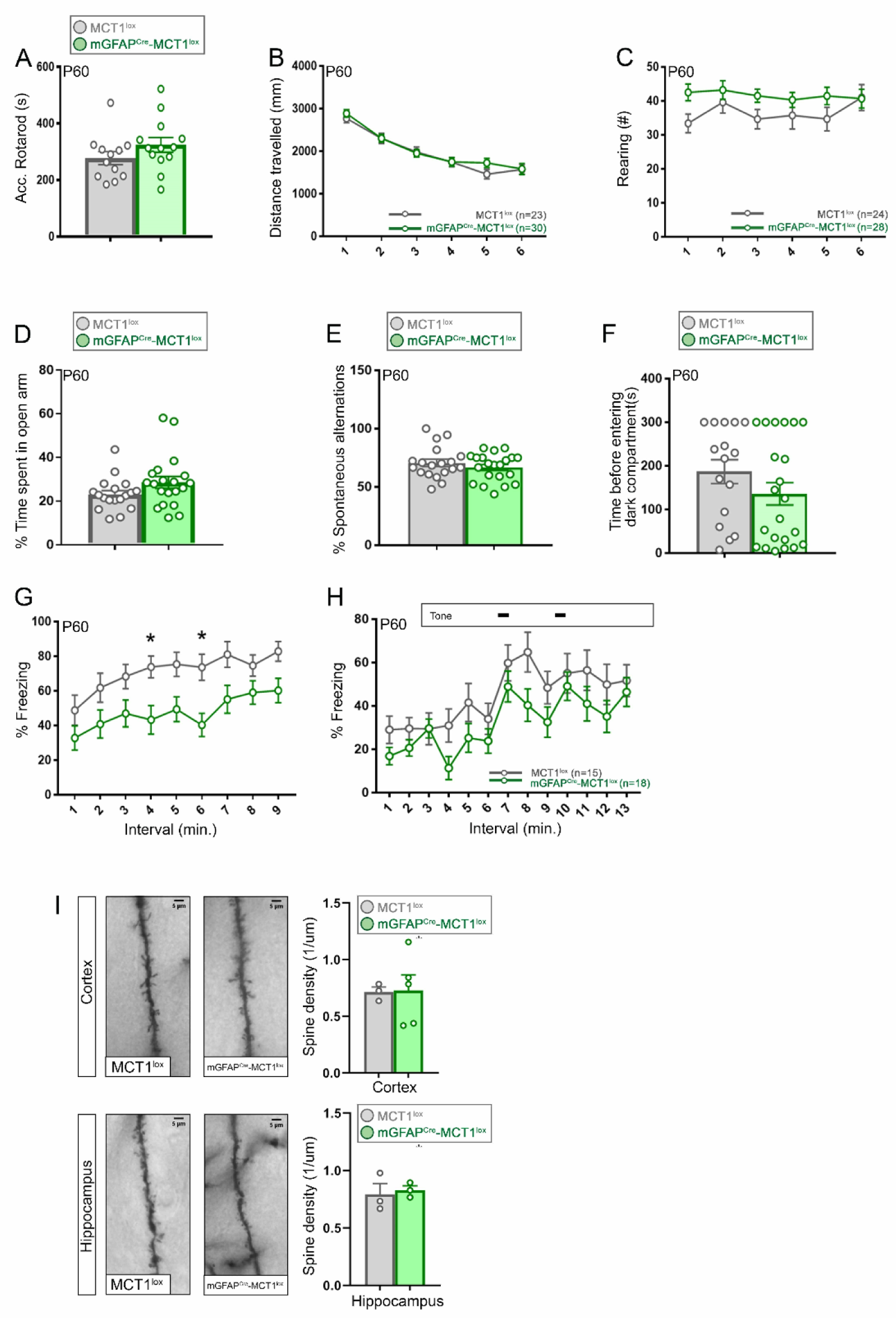
Loss of astrocyte MCT1 does not cause behavioral deficits. (A) P60 mGFAP^Cre^-MCT1^lox^ and MCT1^lox^ controls did not show differences in performance on accelerated rotarod (n=12-13). Data is represented as mean ± SEM. (B) There were no differences between P60 mGFAP^Cre^-MCT1^lox^ and MCT1^lox^ controls for distance traveled during open field testing (n=23-30). Data is represented as mean ± SEM. (C) There were no differences between P60 mGFAP^Cre^-MCT1^lox^ and MCT1^lox^ in rearing activity during open field testing (n=23-30). Data is represented as mean ± SEM. (D) P60 mGFAP^Cre^-MCT1^lox^ and MCT1^lox^ controls spent equal amounts of time in the open arm as measured by elevated plus maze (n=19-20). Data is represented as mean ± SEM. (E) There were no differences in the percentage of spontaneous alternations between P60 mGFAP^Cre^-MCT1^lox^ and MCT1^lox^ mice (n=18-21). Data is represented as mean ± SEM. (F) There were no differences in the amount of time before entering the dark compartment as measured by the passive avoidance test between P60 mGFAP^Cre^-MCT1^lox^ and MCT1^lox^ mice (n=16-22). Data is represented as mean ± SEM. (G). Freezing behavior measured during the context phase of the fear conditioning test was significant reduced at 4 and 6 minutes for mGFAP^Cre^-MCT1^lox^ mice compared to MCT1^lox^ littermate controls. (n=15-18, *p<0.05, two-way ANOVA with Sidak’s multiple comparison’s test) (I) There were no differences in freezing behavior when analyzing P60 mGFAP^Cre^-MCT1^lox^ and MCT1^lox^ mice during the cued phase of the fear trace conditioning test (n=15-18). Data is represented as mean ± SEM. (J) Spine density analysis with Golgi-Cox staining did not reveal differences in either cortex or hippocampus of P360 mGFAP^Cre^-MCT1^lox^ and MCT1^lox^ littermate controls (n=3-4). Scale bar is 5µm. Data is represented as mean ± SEM.

### 3.3 Loss of astrocyte MCT1 is not associated with axonal degeneration during advanced aging

We next determined whether the mGFAP^Cre^-MCT1^lox^ mice develop any signs of neuropathology in different CNS areas with aging. We performed immunofluorescence for neuronal, microglial and astrocytic markers with aging. In addition, we performed transmission electron microscopic analysis of cross sections of optic nerves isolated from mGFAP^Cre^-MCT1^lox^ mice and compared to MCT1^lox^ controls. We did not observe changes in neuronal cell counts, microglial or astrocytic reactivity in either lumbar spinal cord or brain sections from P30 mGFAP^Cre^-MCT1^lox^ mice as compared to controls (Supplementary Figure 3A-B) suggesting that loss of astrocyte MCT1 does not impact normal CNS development. Mice were then aged to P720 and immunostaining was performed on tissue sections from different CNS areas. To our surprise, even at P720, we did not observe changes in astrocyte or microglial reactivity reflective of neuronal degeneration (Figure 3A). We counted neurons and motor neurons in the spinal cord of P360 mGFAP^Cre^-MCT1^lox^ mice and found no difference in the number of NeuN^+^ neuronal or Chat^+^ motor neuron cell bodies (Supplementary Figure 3B-C). We performed electron microscopic analysis of optic nerves isolated from mGFAP^Cre^-MCT1^lox^ and MCT1^lox^ controls. We did not observe overt axonal degeneration in either P360 or P720 optic nerves. There was no change in glial reactivity, or changes in mitochondrial accumulation or density (Figure 3B and C and data not shown). There was no change in glial reactivity in cortex or subcortical areas or indication of neuronal injury in the aged mGFAP^Cre^-MCT1^lox^ mice (Supplementary Figure 3D-E). Myelin thickness analysis in the mGFAP^Cre^-MCT1^lox^ mice did not reveal significant differences at P360 orP720 (Figure 3B). To exclude potential effects of genetic background on the lack of phenotype observed, we backcrossed our B6/SJL conditional nulls to C57Bl6/J mice for > 10 generations. Genetic background did not affect the phenotype observed as axonal degeneration was not observed in C57Bl6/J backcrossed animals at either P180 or P720 (Supplementary Figure 3F).

**Figure 3:**
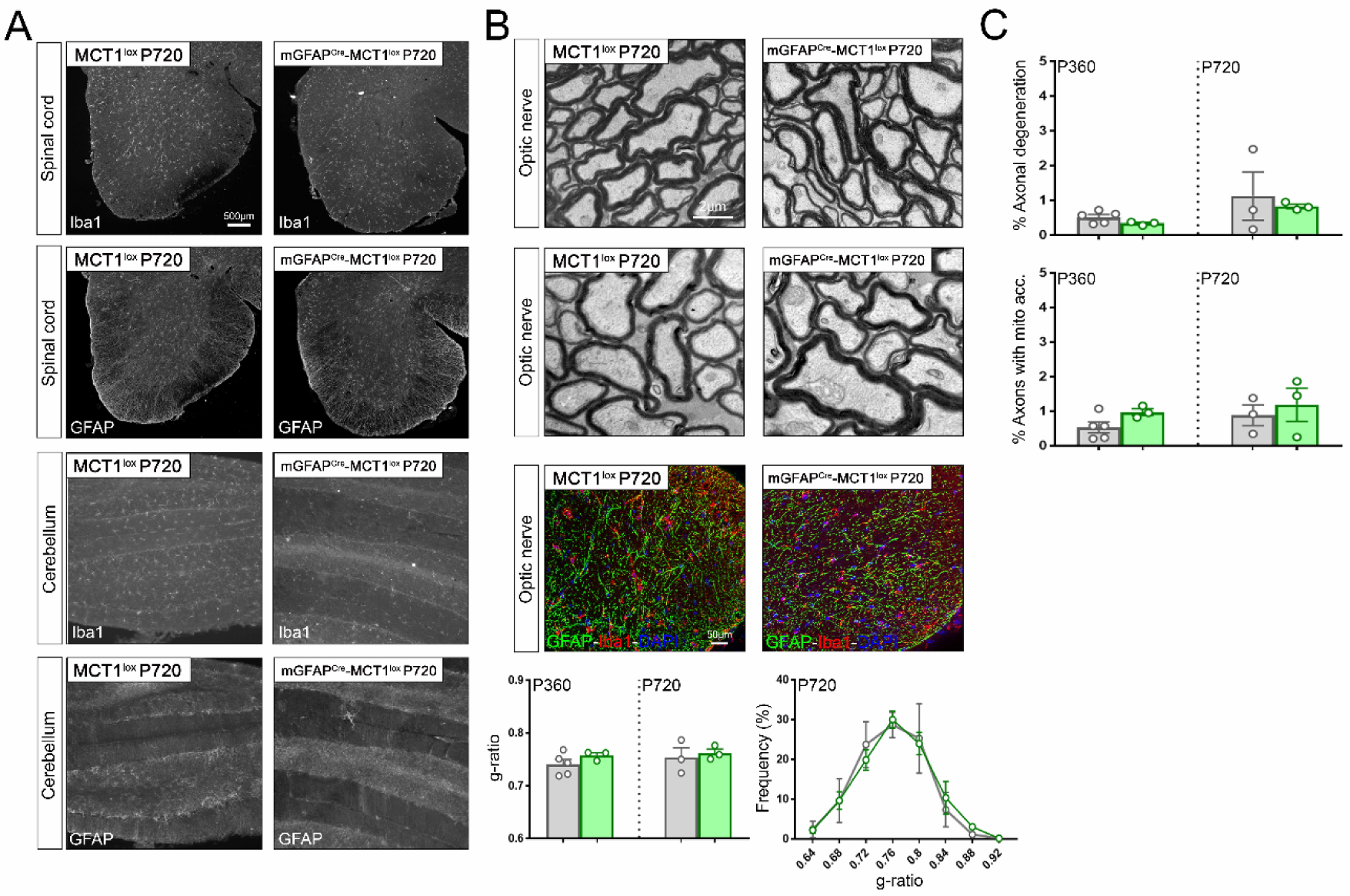
Astrocyte conditional null mice do not develop pathological changes in the CNS with aging. (A) Immunofluorescence from spinal cord and cerebellar sections stained for astrocytes (GFAP) and microglia (Iba1). No increase in staining intensity was observed in the CNS of mGFAP^Cre^-MCT1^lox^ mice as compared to MCT1^lox^ littermate controls. Scale bar is 500µm. Images are representative of n=3 per group. (B) Analysis of optic nerves isolated from P720 mGFAP^Cre^-MCT1^lox^ and MCT1^lox^ littermate control mice. Electron microscopic images did not reveal signs of active axonal degeneration (top panels). Scale bar is 2µm. Immunostaining for astrocyte (GFAP) and microglia (Iba1) does not reveal changes in immunoreactivity (middle panels). Scale bar is 50µm. Quantification of g-ratio and g-ratio frequency distribution does not indicate there are differences between P360 or P720 mGFAP^Cre^-MCT1^lox^ and MCT1^lox^ optic nerves. Data is represented as mean ± SEM. (C) Quantification of top panel images in (B). There was no difference in percentage of axonal degeneration or in the percentage of axons with mitochondrial swellings when comparing P720 mGFAP^Cre^-MCT1^lox^ and MCT1^lox^ mice. Data is represented as mean ± SEM.

### 3.4 Endothelial MCT1 is dispensable for long-term neuronal energy homeostasis

Our conditional MCT1^lox^ mice were crossed with Tie2^Cre^-MCT1^lox^ mice, heterozygous for Tie2 Cre expression and homozygous for the MCT1^lox^ allele. The Tie2 promoter drives Cre expression in endothelial cells and therefore by crossing with the MCT1 conditional null mice we obtain loss of MCT1 expression specifically in endothelial cells from embryogenesis onward. We performed quantitative PCR, Western Blot and immunofluorescence to assess MCT1 knockdown efficiency in endothelial cells. Although we did not obtain a significant reduction in *Mct1* mRNA in the different CNS areas analyzed, we did observe a 35% reduction in MCT1 protein in spinal cord whereas the reduction in MCT1 protein expression in the cortex did not reveal a significant difference (Supplementary Figure 4A-B). The lack of significant downregulation is likely due to high MCT1 expression in unaffected cell types in this conditional knockout approach, e.g. MCT1 expression in oligodendrocytes and astrocytes. In fact, when assessing MCT1 expression by immunostaining on lumbar spinal cord cross sections of P30 Tie2^Cre^-MCT1^lox^ and MCT1^lox^ littermate controls, we observed a near complete loss of MCT1 expression in endothelial cells, validating our conditional knockout approach (Supplementary Figure 4C). To assess whether loss of endothelial cell MCT1 leads to vascular leakage, we injected tomato Dylight 488 Lectin intracardially into anesthetized mice. Five minutes after injection, mice were euthanized, and brain sections were cut and visualized with fluorescent microscopy. We did not observe leakage of the blood vasculature as all DyLight labeling was still associated with blood vessels in both the Tie2^Cre^-MCT1^lox^ as well as MCT1^lox^ aged-matched controls (Supplementary Figure 4D). Lastly, we performed immunofluorescent and electron microscopic analysis of different CNS areas looking for any signs of axonal pathology or neuronal degeneration. Neither immunostaining nor electron microscopic analysis revealed any differences in microglial or astrocytic reactivity in the cortex, spinal cord, hippocampus or cerebellum, or the presence of degenerating axons in the optic nerve axons even in mice aged to P720. In summary, these data indicate that loss of MCT1 from endothelial cells, similarto what is observed upon astrocytic conditional deletion, is dispensable for endothelial function and the maintenance of neuronal metabolic integrity with aging.

### 3.5 Ubiquitous loss of MCT1 causes widespread axonal degeneration

As there was no obvious phenotype observed upon conditional loss of MCT1 expression in either astrocytes or endothelial cells, we reasoned that different cell subtypes expressing MCT1 mediate neuronal metabolic support and potentially compensate for the loss of MCT1 in a solitary cell subtype. We therefore used an approach in which MCT1 is reduced more ubiquitously that would avoid issues arising from full MCT1 deletion, which is embryonic lethal. We crossed CAGG^CreER^ mice with conditional MCT1^lox^ mice and analyzed mice heterozygous for CAGG^CreER^ and homozygous for the MCT1^lox^ allele. To assess the efficiency of MCT1 knockdown in the CAGG^CreER^-MCT1^lox^ mice, CAGG^CreER^-MCT1^lox^ and MCT1^lox^ mice were injected with tamoxifen at P60 and euthanized 4-6 months later to compare MCT1 expression by Western Blot, immunofluorescence and qPCR. MCT1 protein expression was reduced by 61% in the cortex and by 82% in the spinal cord 6 months after tamoxifen injection (Figure 4A). *Mct1* mRNA expression was reduced by 93% in the cortex and by 83% in the spinal cord (Supplementary Figure 5A).

**Figure 4:**
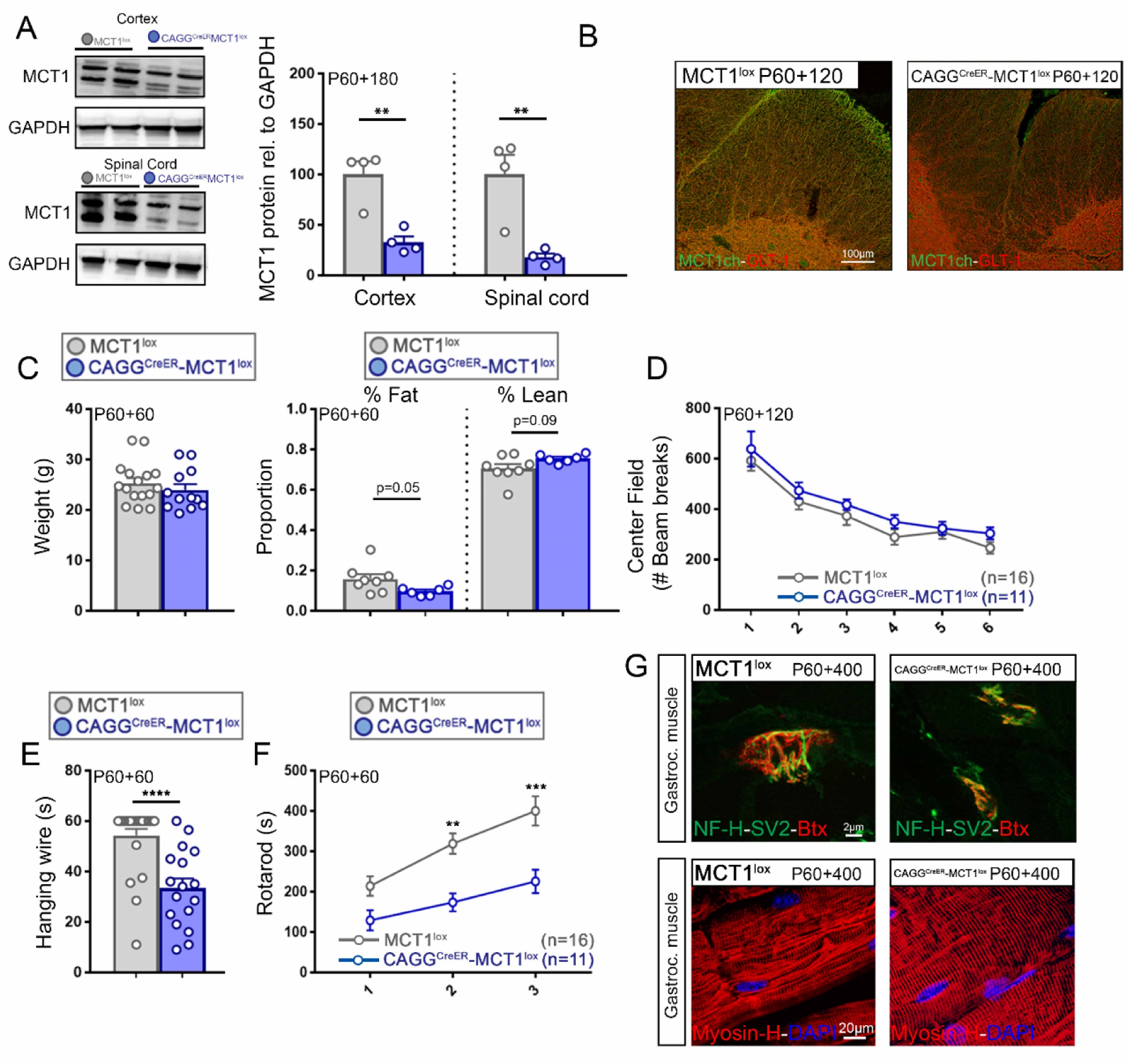
Loss of MCT1 and behavioral changes in ubiquitous MCT1 conditional null mice. (A) MCT1 expression as measured by Western Blot reveals a 61% reduction of MCT1 protein in the cortex of P60+180 CAGG^CreER^MCT1^lox^ mice as compared to MCT1^lox^ controls (n=4, **p<0.01, Student’st-test), whereas spinal cord MCT1 expression was reduced by 82% (n=4, **p<0.01, Student’s t-test). Data is represented as mean ± SEM. (B) Immunostaining of spinal cords from P60+120 CAGG^CreER^-MCT1^lox^ mice reveal a striking loss of MCT1 expression as compared to MCT1^lox^ mice. Note the loss of MCT1 on the GLT-1 expressing astrocyte processes. Scale bar is 100µm. Images are representative of a minimum of n=3 per group. (C) At P60+60 there was no difference in the weight or the percentage fat/lean between CAGG^CreER^MCT1^lox^ and MCT1^lox^ mice (n=12-16). Data is represented as mean ± SEM. (D) There were no differences in the amount of time spent in the center of the field as measured by open field testing (n=11-16). Data is represented as mean ± SEM. (E) There was a 50% reduction in hanging wire performance in the CAGG^CreER^MCT1^lox^ mice as compared to the MCT1^lox^ littermate controls (n=16-17, ****p<0.0001). Data is represented as mean ± SEM. (F) There was a significant loss in accelerated rotarod performance in the CAGG^CreER^-MCT1^lox^ mice as compared to the MCT1^lox^ littermate controls (n=11-16, **p<0.01, ***p<0.001). Data is represented as mean ± SEM. (G) There were no differences in muscular innervation (top panels) or gastrocnemius Myosin-H immunolabeling between the CAGG^CreER^-MCT1^lox^ and MCT1^lox^ mice as compared to the MCT1lox littermate controls. Scale bar is 5µm in upper panels and 20µm in lower panels. Images are representative of a minimum of n=3 per group

These findings were confirmed by immunofluorescence of white and grey matter spinal cord where MCT1 expression was strongly decreased in endothelial cells, as well as, GLT-1 expressing astrocyte processes in the CAGG^CreER^-MCT1^lox^ mice in contrast to the expression observed in MCT1^lox^ controls (Figure 4B). We did not obtain evidence for alterations in expression of other MCTs, connexins or glucose transporters when comparing CAGG^CreER^-MCT1^lox^ and MCT1^lox^ mice that were injected with tamoxifen at P30 and euthanized 30 days later (P30+30) (Supplementary Figure 5B-D). We then injected CAGG^CreER^-MCT1^lox^ and MCT1^lox^ animals with tamoxifen at P60 and analyzed animals 2 months later with a battery of behavioral tests. There were no differences in body weight between CAGG^CreER^-MCT1^lox^ and MCT1^lox^ mice, but there was a trend toward differences in % body fat and lean weight when comparing both genotypes with echo-magnetic resonance imaging (Figure 4C). General locomotion activity was not affected in CAGG^CreER^-MCT1^lox^ mice as compared to MCT1^lox^ controls, but CAGG^CreER^-MCT1^lox^ mice displayed significant decreased ability on the hanging wire test and on accelerated rotarod (Figure 4E-F and Supplementary Figure 5E-G). To rule out effects on muscular innervation and muscular activity, we performed immunostaining for neuronal markers NF-H and SV-2 and postsynaptic nicotinic receptors with α-bungarotoxin but did not see evidence of muscular denervation (Figure 4G). In addition, myosin-H stained gastrocnemius muscle did not reveal signs of muscular degeneration (Figure 4G). Subsequent evaluation of CAGG^CreER^-MCT1^lox^ and MCT1^lox^ mice with behavior studies for anxiety (Supplementary Figure 5G-H) and memory (Supplementary Figure 5I-K) did not reveal changes between both genotypes, suggesting that apart from a prominent loss of motor function, ubiquitous conditional loss of MCT1 expression at adult age (P60) is well tolerated with aging.

We then performed a pathological examination of different CNS regions of CAGG^CreER^-MCT1^lox^ and MCT1^lox^ mice. Within the first three months after tamoxifen injection (P60+100) we did not observe any signs of ongoing neurodegeneration such as glial reactivity (Supplementary Figure 6A) and therefore opted to age the CAGG^CreER^-MCT1^lox^ and MCT1^lox^ mice to 400 days post tamoxifen injection (P60+400) and performed immunofluorescence for glial cell reactivity markers and electron microscopy for signs of axonal degeneration. Interestingly, we observed localized increases in glial reactivity in the brain stem and cerebellum (Figure 5A and Supplementary Figure 6C). In the spinal cord we observed a trend towards enhanced astrocytic reactivity in the grey matter area, but we did not observe differences in the number of NeuN^+^ neurons between CAGG^CreER^-MCT1^lox^ and MCT1^lox^ controls (Figure 5B and Supplementary Figure 6B). We also performed electron microscopy evaluation of the dorsal corticospinal tract but did not observe any signs of axonal degeneration (Figure 5B). Interestingly, we observed profound axonal degeneration in the optic nerves isolated from the CAGG^CreER^-MCT1^lox^ mice but not the MCT1^lox^ aged-matched controls, which correlated with an increase in both astrocytic and microglial reactivity in this area (Figure 5C-F). We observed dark, degenerated axons and axons with swollen cytoplasm and accumulated intracellular material undergoing active degeneration. In P60+400 CAGG^CreER^-MCT1^lox^ mice, mitochondrial size was unaffected, though there was a trend towards an increase in the percentage of axons that had accumulated multiple mitochondria, which could be reflective of underlying metabolic defects (Figure 5F-G). Interestingly, the percentage of degenerated axons in P60+400 CAGG^CreER^-MCT1^lox^ mice was significantly increased as compared to MCT1^lox^ littermate controls (Figure 5H).

**Figure 5:**
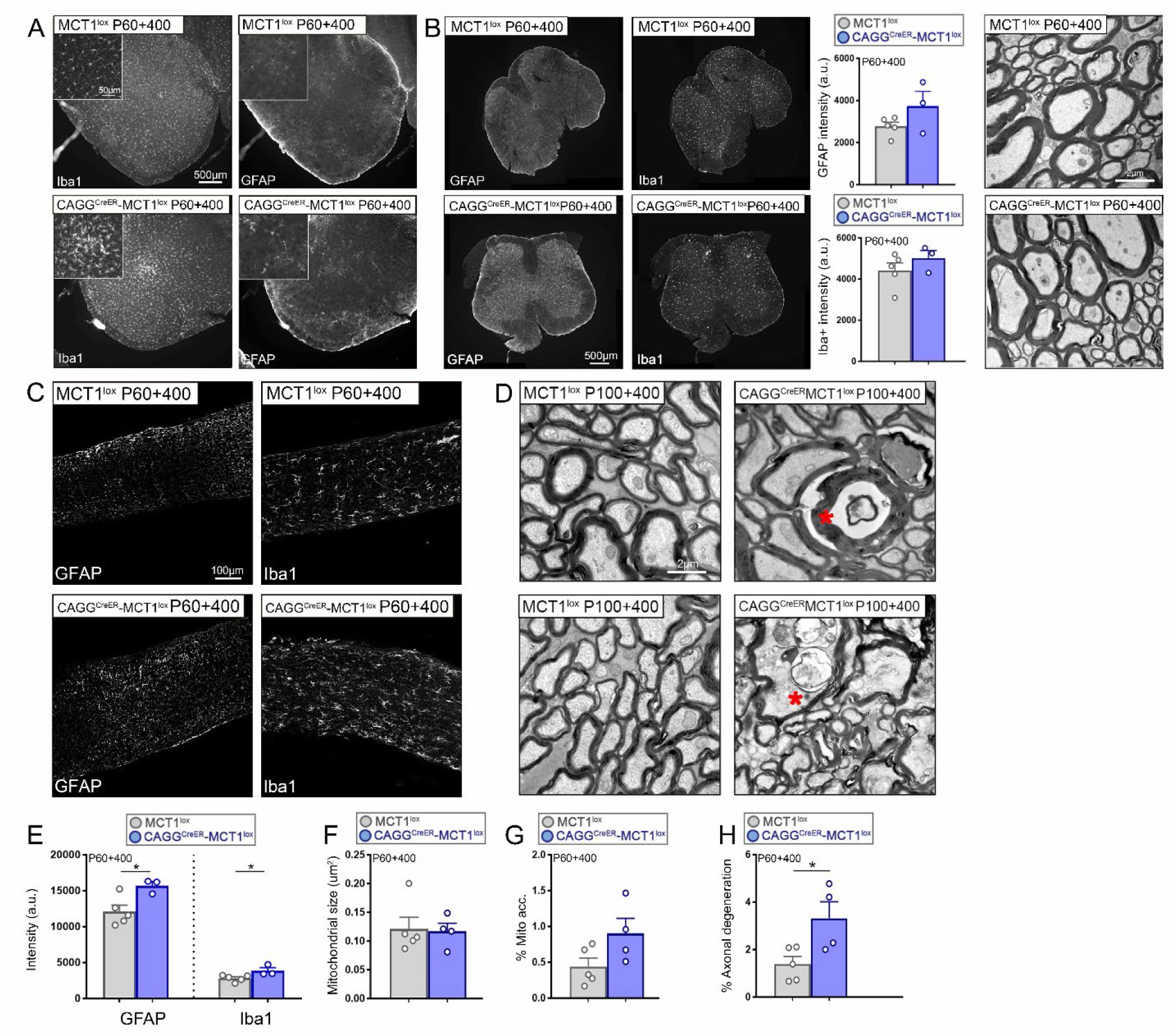
Ubiquitous loss of MCT1 causes widespread axonal degeneration. (A) Immunofluorescence of brain stem reveals patches of enhanced immunoreactivity for microglial marker Iba1 in mGFAP^Cre^-MCT1^lox^ as compared to MCT1^lox^ controls. Scale bar is 50µm in small panels and 500µm in large panels. Images are representative of a minimum of n=3 per group (B) Immunofluorescence of lumbar spinal cord for astrocyte (GFAP) and microglia (Iba1) markers reveals a lack of overall increase in marker expression in CAGG^CreER^-MCT1^lox^ as compared to MCT1^lox^ controls. Electron microscopic evaluation from the spinal cord dorsal corticospinal tract (panels on the right) does not reveal signs of ongoing axonal degeneration. Images are representative of a minimum of n=3 per group. Scale bar is 500µm in immunofluorescent images and 2µm in EM images. Data is represented as mean ± SEM. (C) Immunofluorescence of optic nerves for astrocyte (GFAP) and microglia (Iba1) markers reveals an increase of overall marker expression in P60+400 CAGG^CreER^-MCT1^lox^ as compared to age-matched MCT1^lox^ controls. Scale bar is 100µm. (D) Electron microscopic evaluation of optic nerves dissected from CAGG^CreER^-MCT1^lox^ and MCT1^lox^ controls. Note the presence of active axonal degeneration with accumulation of intracellular organelles (red asterisk). Scale bar is 2µm. (E) Quantification of immunofluorescence images from (C) (n=3-5, *p<0.05, Student’s t-test). Data is represented as mean ± SEM. (F-H) Quantification of mitochondrial size, mitochondrial accumulation, and axonal degeneration in CAGG^CreER^-MCT1^lox^ mice as compared to MCT1^lox^ controls. (n=4-5, *p<0.05, Student’s t-test).

## 4. Discussion

Astrocytes support neurons through a multitude of different actions among which maintenance of neuronal energy metabolism, regulation of glutamate recycling and antioxidant protection have been extensively studied. One of the key pathways that provides metabolic support to neurons is the so-called astrocyte-neuron lactate shuttle, in which astrocytes produce lactate in response to neuronal activity and shuttle lactate to neurons to fuel neuronal metabolic demands(Pellerin and Magistretti, 1994, Philips and Rothstein, 2017). Numerous studies have hinted that astrocyte-derived lactate is essential for neuronal metabolic support(Descalzi et al., 2019, Tekkok et al., 2005, Vezzoli et al., 2019, Suzuki et al., 2011, Netzahualcoyotzi and Pellerin, 2020). Extracellular glucose and lactate concentrations when measured with bio-probes or biosensors change as mice perform a memory task suggesting a role for lactate during memory formation(Newman et al., 2011). Interestingly, inhibition of glycogenolysis or downregulation of monocarboxylate transporters MCT1 or MCT4leads to long-term potentiation (LTP) impairment and amnesia in an inhibition/avoidance (I/A) learning task in rats, which could be rescued by the administration of l-lactate(Suzuki et al., 2011). In the mice hippocampus, I/A learning leads to an upregulation in the expression of several metabolic genes including MCTs, glucose transporter 1 and the alpha2 subunit of the Na/K-ATPase(Tadi et al., 2015). Loss of MCT1 expression in the MCT1 heterozygous null mice impaired memory and performance in the I/A test(Tadi et al., 2015). Similarly, when mice were tasked to learn a novel object recognition task, the resulting long-term memory formation was associated with an increase in spine density and enhanced astrocytic glycogen granule density(Vezzoli et al., 2019). Interestingly, inhibition of glycogen metabolism by DAB prevented these synaptic changes and mice were unable to learn the memory task. Again, injecting lactate into the hippocampus rescued this phenotype, suggesting that glycogen-derived lactate has an important role in learning-induced synaptic plasticity. With this in mind, we were surprised to observe that the conditional deletion of MCT1 from astrocytes did not show a similar phenotype to those observed in these earlier studies. We did not observe changes in spine density in the hippocampus, nor did we observe changes in memory tasks, including I/A testing, Y-maze test and fear conditioning. In fact, a whole battery of behavioral and pathological examinations failed to reveal any sign of ongoing degeneration suggesting that astrocytic MCT1 is dispensable for long-term maintenance of axonal energy homeostasis. Multiple reasons could be given for these discrepancies. Firstly, we have a model with chronic inhibition of MCT1 in astrocytes that begins during development, which could induce compensatory metabolic mechanisms that bypass the need for MCT1 transporters in e.g. memory establishment. We did perform an analysis of the expression of alternative metabolic pathways like glucose transporters and connexin hemichannels and although we failed to observe changes in expression, this does not rule out the involvement of an alternative route of axonal energy homeostasis during the lifespan. For example, it has been shown that astrocytes can accumulate lactate at concentrations above thermodynamic equilibrium and 37 pS lactate-permeable ion channels expressed by astrocytes can cause fast lactate depletion in response to changes in extracellular potassium (Sotelo-Hitschfeld et al., 2015). Secondly, MCT1 is expressed by multiple, if not all, cell types in the CNS, and therefore multiple cell types could compensate for the loss of astrocytic MCT1 function. We recently reported that mice with conditional deletion of MCT1 in oligodendrocyte lineage cells develop axonal degeneration by one year of age. These mice developed normally without overt behavioral changes, but by one year of age these animals acquired a subtle axonal degeneration phenotype in optic nerve and patchy glial cell activation in different CNS areas. With subsequent aging to 2 years of age, these mice have even more pronounced degenerative phenotypes observed upon electron microscopic pathological evaluation of the optic nerves. In contrast, our studies with conditional astrocytic MCT1 deletion did not lead to animal behavioral changes or overt degenerative phenotypes even with advanced aging. This discrepancy with the previously reported learning/memory deficiencies could well be due to the fact that these studies lack cell-specific targeting, e.g. by injection of MCT1 ASOs that do not have cell-specific targeting capabilities. Lastly, our behavioral experiments were performed in a mixed B6/SJL background--as changes in genetic background do have particular effects on mouse metabolism, this could explain differences to what is observed with I/A testing in heterozygous MCT1 null mice with B6 background.

Similar to our observations with the astrocyte conditional null mice, we did not observe pathology in the endothelial conditional null mice, even when aged to 2 years. As MCT1 is involved in angiogenesis in glioblastoma, we wondered whether loss of endothelial MCT1 would induce an aggressive phenotype. However, vascularization was indistinguishable in the Tie2^Cre^-MCT1^lox^ mice when compared to their age-matched MCT1^lox^ controls. The lack of axonopathy might not be that surprising given that endothelial cells lack the intricate interaction with axons like oligodendrocytes (through their myelin sheath) and astrocytes (through the tripartite synapse) have. Nevertheless, endothelial MCT1 has been shown to control extracellular lactate levels that impact hippocampal neurogenesis. In addition, endothelial derived lactate controls M2 polarization in peripheral macrophages and muscle regeneration upon ischemia which depends on endothelial MCT1 expression. We conclude for our studies that under normal homeostatic conditions endothelial MCT1 is dispensable, although we can not rule out milder underlying changes in axonal or glial homeostasis that do not lead to an overt phenotype. In addition, more in-depth analysis of these endothelial conditional null mice could shed light on the role of endothelial MCT1 in CNS neuro-injury models.

As phenotypes are rather mild or entirely absent upon conditional depletion of MCT1 from particular glial cells, we wondered whether a ubiquitous approach would lead to more pronounced metabolic disturbances and degeneration in axons throughout the central nervous system. We used the CAGG^CreER^ mouse, which not only allows for the conditional depletion of MCT1 throughout the entire but also recombines the floxed MCT1 allele in peripheral organs. In addition, it allows for the assessment of MCT1 importance in neuronal energy homeostasis after the mouse has developed normally, to rule out any potential toxic effect that occurs during development as observed in full MCT1 null mice who die embryonically. Interestingly, we did observe profound axonal degeneration upon ubiquitous conditional loss of MCT1 about one year after tamoxifen injection, particularly in the mouse optic nerve. No such degeneration was observed at similar or later time points in mice with conditional depletion of MCT1 specifically in either astrocytes or endothelial cells.

This study highlights that chronic depletion of MCT1 from astrocytes or endothelial cells is well tolerated during mouse development as well as with aging up to 2 years of age. Ubiquitous depletion of MCT1 using conditional null mice lead to phenotypes similar to those obtained with the oligodendrocyte conditional null mice as reported earlier. We therefore conclude that unlike oligodendrocyte MCT1, loss of astrocyte MCT1 is well tolerated with aging and astrocyte MCT1 is not an essential driver of astrocyte mediated axonal energy homeostasis with aging.

## Acknowledgements

This work was funded by the Muscular Dystrophy Association (MDA, development grant 381190) (T.P.), ALS Association (J.D.R.), Department of Defense (J.D.R.) and NIH (J.D.R.). We acknowledge Carol Cooke for her assistance with the EM at the Neurology-Peripheral Nerve Division at Johns Hopkins University School of Medicine. We thank members of J.D.R. lab at Johns Hopkins University School of Medicine for helpful discussions.

---

**Supplementary Figure 1**

(A) Immunofluorescence staining for MCT1 in spinal cord (upper panels) and cortex (lower panels) of mGFAP^Cre^-MCT1^lox^ and MCT1^lox^ controls. Scale bar is 100µm. Images are representative of n=3 per group.

(B) *Mct1* mRNA expression in the spinal cord of mGFAP^Cre^-MCT1^lox^ mice was reduced by 24% as compared to MCT1^lox^ controls, whereas mRNA expression levels for *Mct2* and *Mct4* was unaffected (n=6-8, ***p<0.001, Student’s t-test). Scale bar is 200µm. Data is represented as mean ± SEM

(C) There were no changes in mRNA expression levels of connexin hemichannels *Cx43, Cx30* and *Cx47* or glucose transporter subtypes *Glut1* and *Glut3* between mGFAP^Cre^-MCT1^lox^ and MCT1^lox^ mice (n=3-5). Data is represented as mean ± SEM.

(D) There were no changes in the expression of MCT1 in the myelin derived from whole spinal cord of P30 mGFAP^Cre^-MCT1^lox^ as compared to age-matched MCT1^lox^ mice (n=3). The MCT1 protein expression in the ‘total’ fraction (entire spinal cord after removal of myelin) was significantly reduced in the spinal cord of mGFAP^Cre^-MCT1^lox^ as compared to MCT1^lox^ mice (n=3, *p<0.05, Student’s t-test). Data is represented as mean ± SEM.

**Supplementary Figure 2**

(A) There was no significant difference in weight between P60 mGFAP^Cre^-MCT1^lox^ and MCT1^lox^ mice (n=7-11). Data is represented as mean ± SEM.

(B) There were no significant changes in fat and lean mass measured by echo-MRI when comparing P60 mGFAP^Cre^-MCT1^lox^ and MCT1^lox^ mice (n=8-11). Data is represented as mean ± SEM.

(C) The number of arm entries during Y-maze testing was similar between P60 mGFAP^Cre^-MCT1^lox^ and MCT1^lox^ mice (n=19-20). Data is represented as mean ± SEM.

(D) Percentage of time the animals freeze during the preconditioning stage of the fear trace conditioning test in mGFAP^Cre^-MCT1^lox^ and MCT1^lox^ mice (n=15-18). Data is represented as mean ± SEM.

(E) Percentage of time the animals freeze during the training stage of the fear trace conditioning test of mGFAP^Cre-^MCT1^lox^ and MCT1^lox^ mice (n=15-18). Data is represented as mean ± SEM.

(F) There are no differences in NeuN^+^ neuronal cell count in the CA1 region of the hippocampus when comparing P720 old mGFAP^Cre^-MCT1^lox^ with age-matched MCT1^lox^ controls (n=5-7). Data is represented as mean ± SEM.

**Supplementary Figure 3**

(A) Immunostaining of astrocyte (GFAP) and microglia (Iba1) markers in P30 spinal cord and neuronal marker NeuN in P90 coronal brain sections of mGFAP^Cre^-MCT1^lox^ and MCT1^lox^ mice. Quantification of NeuN^+^ cells in the different cortical layers did not reveal differences in NeuN^+^ cell number (n=4). Scale bar is 200µm. Data is represented as mean ± SEM.

(B) NeuN immunostaining in spinal cord of P30 and P360 mGFAP^Cre^-MCT1^lox^ and MCT1^lox^ mice. Quantification of NeuN+ cells did not reveal changes at both time points analyzed (n=3-7). Scale bar is 100µm. Data is represented as mean ± SEM.

(C) Chat immunostaining in spinal cord of P360 mGFAP^Cre^-MCT1^lox^ and MCT1^lox^ mice. Quantification of Chat^+^ cells did not reveal changes at both time points analyzed (n=3-7). Scale bar is 100µm. Data is represented as mean ± SEM.

(D) Immunostaining of astrocyte (GFAP) and microglia (Iba1) markers in coronal brain section of P360 mGFAP^Cre^-MCT1^lox^ and MCT1^lox^ mice. Scale bar is 200µm. Images are representative of n=3.

(E) Electron microscopy images of the cortex and hippocampus of P360 mGFAP^Cre^-MCT1^lox^ and MCT1^lox^ mice. No abnormalities indicative of ongoing neuronal degeneration was observed in either region. Images are representative of a minimum of n=3. Scale bar is 2µm.

(F) Electron microscopy images of optic nerves isolated from of P180 and P720 mGFAP^Cre^-MCT1^lox^ and MCT1^lox^ mice backcrossed to B6 background for >10 generations. There were no changes in the percentage of accumulating mitochondria or the percentage of axonal degeneration between the two genotypes analyzed (n=3-4). Scale bar is 2µm.

**Supplementary Figure 4**

(A) There were no changes in *Mct1, Mct2* and *Mct4* mRNA expression levels in the spinal cord of P50 Tie2^Cre^-MCT1^lox^ and MCT1^lox^ mice (n=3-6). Data is represented as mean ± SEM.

(B) MCT1 protein expression in the spinal cord of P50 Tie2^Cre^-MCT1^lox^ mice is reduced by 35% when compared to MCT1^lox^ age-matched controls (n=3-4), *p<0.05, Student’s t-test. There were no changes in MCT1 protein expression observed in the cortex. Data is represented as mean ± SEM.

(C) Immunostaining of MCT1 in the spinal cord of P30 MCT1^lox^ mice reveals multiple blood vessels immunoreactive for MCT1, whereas MCT1 immunostaining in Tie2^Cre^-MCT1^lox^ mice is devoid of immunolabeling in blood vessels. Scale bar is 100µm. Images are representative of a minimum of n=3.

(D) Intracardial injected fluorescently labeled lectin was detected throughout the brain of both P30 MCT1^lox^ and Tie2^Cre^-MCT1^lox^ mice suggesting loss of MCT1 in endothelial cells does not cause vascular leakage. Scale bar is 200µm. Images are representative of a minimum of n=3.

(E) Immunostaining of astrocyte (GFAP) and microglia (Iba1) markers in coronal brain sections of P240 Tie2^Cre^-MCT1^lox^ and MCT1^lox^ mice. Scale bar is 500µm. Images are representative of a minimum of n=3.

(F) Electron microscopy images of optic nerves isolated from P720 Tie2^Cre^-MCT1^lox^ and MCT1^lox^ age-matched controls. There were no changes in the percentage of accumulating mitochondria or the percentage of axonal degeneration between the two genotypes analyzed (n=2-3). Scale bar is 2µm.

**Supplementary Figure 5**

(A) The expression of *Mct1* mRNA in the cortex and spinal cord of P30+30 CAGG^CreER^-MCT1^lox^ was reduced by 93 and 89% respectively as compared to MCT1^lox^ controls (n=4), Data is represented as mean ± SEM.

(B) There were no changes in mRNA expression levels of connexin hemichannels *Cx43, Cx30* and *Cx47* between P30+30 CAGG^CreER^-MCT1^lox^ and MCT1^lox^ mice (n=3). Data is represented as mean ± SEM.

(C) There were no changes in mRNA expression levels of glucose transporter subtypes *Glut1* and *Glut3* between P30+30 old CAGG^CreER^-MCT1^lox^ and MCT1^lox^ mice (n=3). Data is represented as mean ± SEM.

(D) Comparison of *Mct1* mRNA expression in the spinal cord of P30+30 old CAGG^CreER^-MCT1^lox^ mice as compared to MCT1^lox^ mice reveals a 83% decrease in expression, whereas mRNA expression levels for *Mct2* and *Mct4* were unaffected (n=3, ***p<0.001, Student’s t-test). Data is represented as mean ± SEM.

(E) There were no differences between P60+120 CAGG^CreER^-MCT1^lox^ and MCT1^lox^ controls for distance traveled as measured by open field testing (n=11-16). Data is represented as mean ± SEM.

(F) There were no differences in average speed during open field testing between P60+120 CAGG^CreER^-MCT1^lox^ and MCT1^lox^ controls (n=11-16). Data is represented as mean ± SEM.

(G) There were no differences in rearing activity during open field testing between P60+120 CAGG^CreER^-MCT1^lox^ and MCT1^lox^ controls (n=11-16). Data is represented as mean ± SEM.

(H) P60+120 old CAGG^CreER^-MCT1^lox^ and MCT1lox mice spent an equal amount of time in the open arms of the elevated plus maze (n=12-17). Data is represented as mean ± SEM.

(I) There were no differences in the amount of time before entering the dark compartment as measured by the passive avoidance test between P60+120 CAGG^CreER^-MCT1^lox^ and MCT1^lox^ mice (n=11-16). Data is represented as mean ± SEM.

(I) There were no differences in the percentage of spontaneous alternations during Y-maze testing between P60+120 CAGG^CreER^-MCT1^lox^ and MCT1^lox^ mice (n=12-17). Data is represented as mean ± SEM.

(I) There were no differences in the number of arm entries during Y-maze testing between P60+120 CAGG^CreER^-MCT1^lox^ and MCT1^lox^ mice (n=12-17). Data is represented as mean ± SEM.

**Supplementary Figure 6**

(A) Immunostaining of astrocyte (GFAP) and microglia (Iba1) markers in spinal cord of P30+70 CAGG^CreER-^MCT1^lox^ and MCT1^lox^ mice. Images are representative of a minimum of n=3. Scale bar is 200µm.

(B) Quantification of NeuN^+^ cells in the spinal cord of P60+400 CAGG^CreER-^MCT1^lox^ and MCT1^lox^ mice did not reveal changes in neuronal cell counts. Images are representative of a minimum of n=3-5. Scale bar is 200µm.

(C) Immunostaining of astrocyte (GFAP) and microglia (Iba1) markers in coronal brain sections of P90+400 CAGG^CreER-^MCT1^lox^ and MCT1^lox^ mice reveals patchy increases in glial reactivity in white matter areas. Images are representative of a minimum of n=3. Scale bar is 100um in small panels and 200µm in large panels.

