## Supplementary figures and images for "Astrocyte MCT1 expression does not contribute to the axonal degenerative phenotype observed with ubiquitous MCT1 depletion"

### Supplemental files

Supp Figure 1

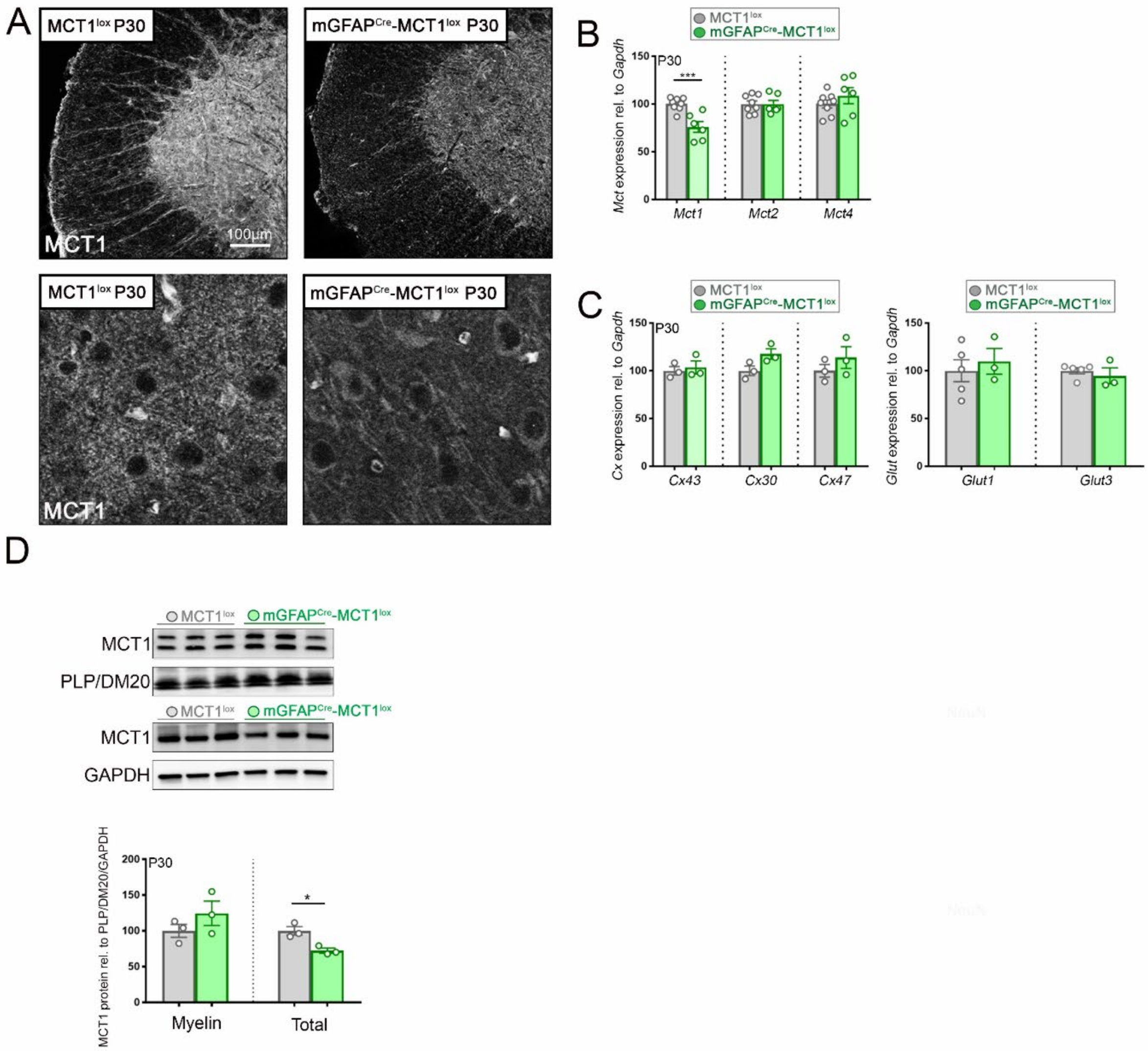

Supp Figure 2

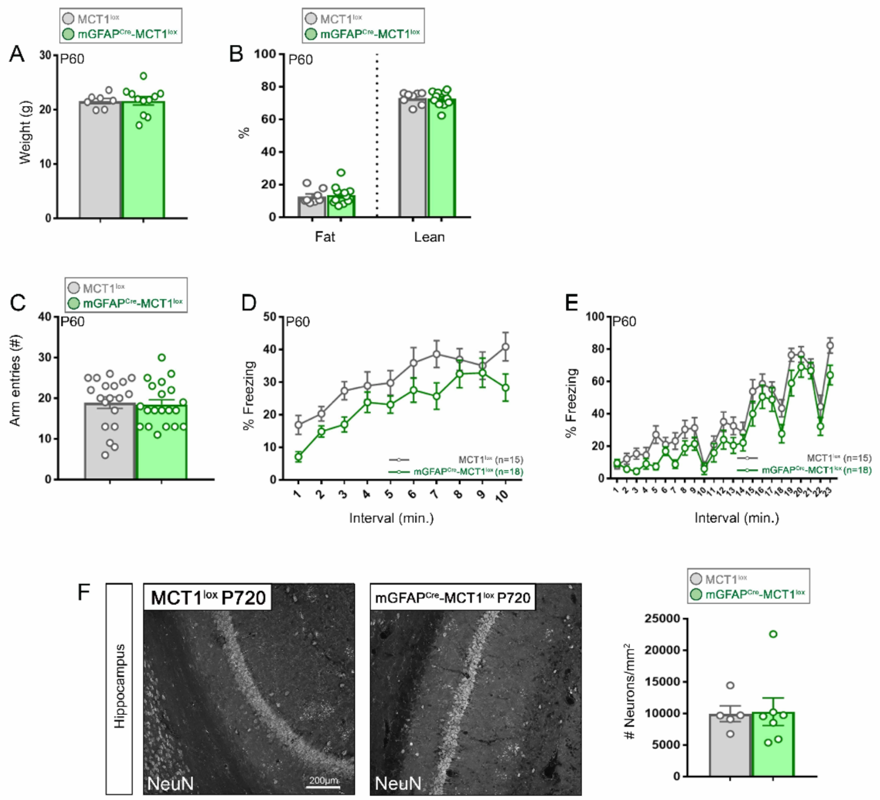

Supp Figure 3

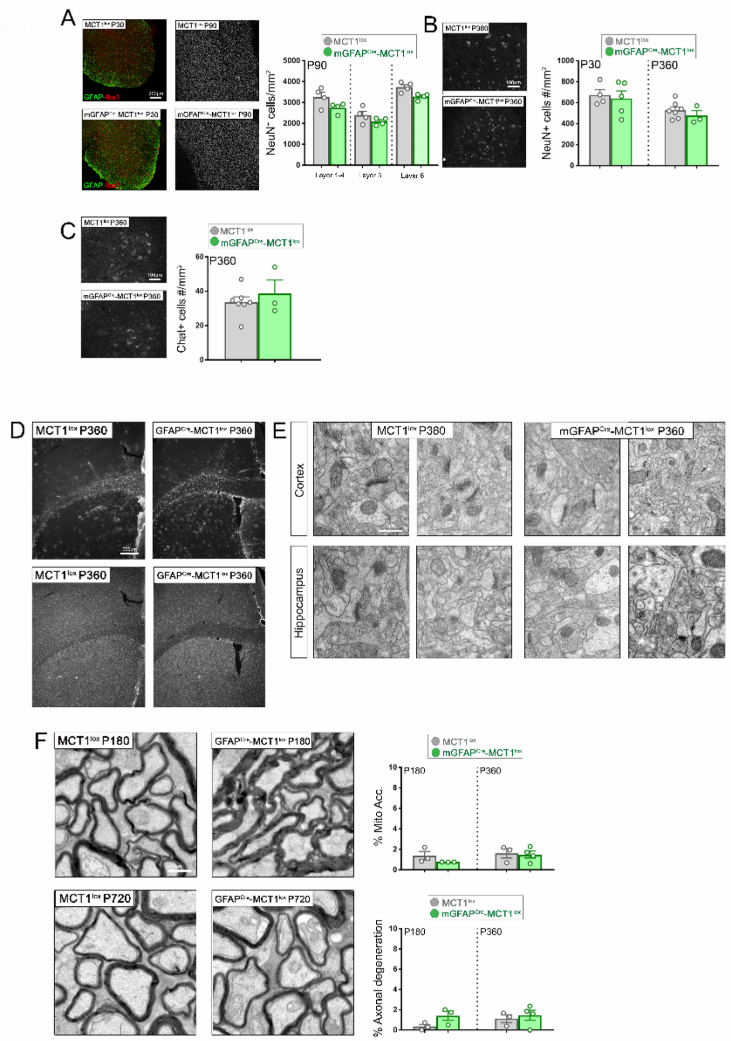

Supp Figure 4

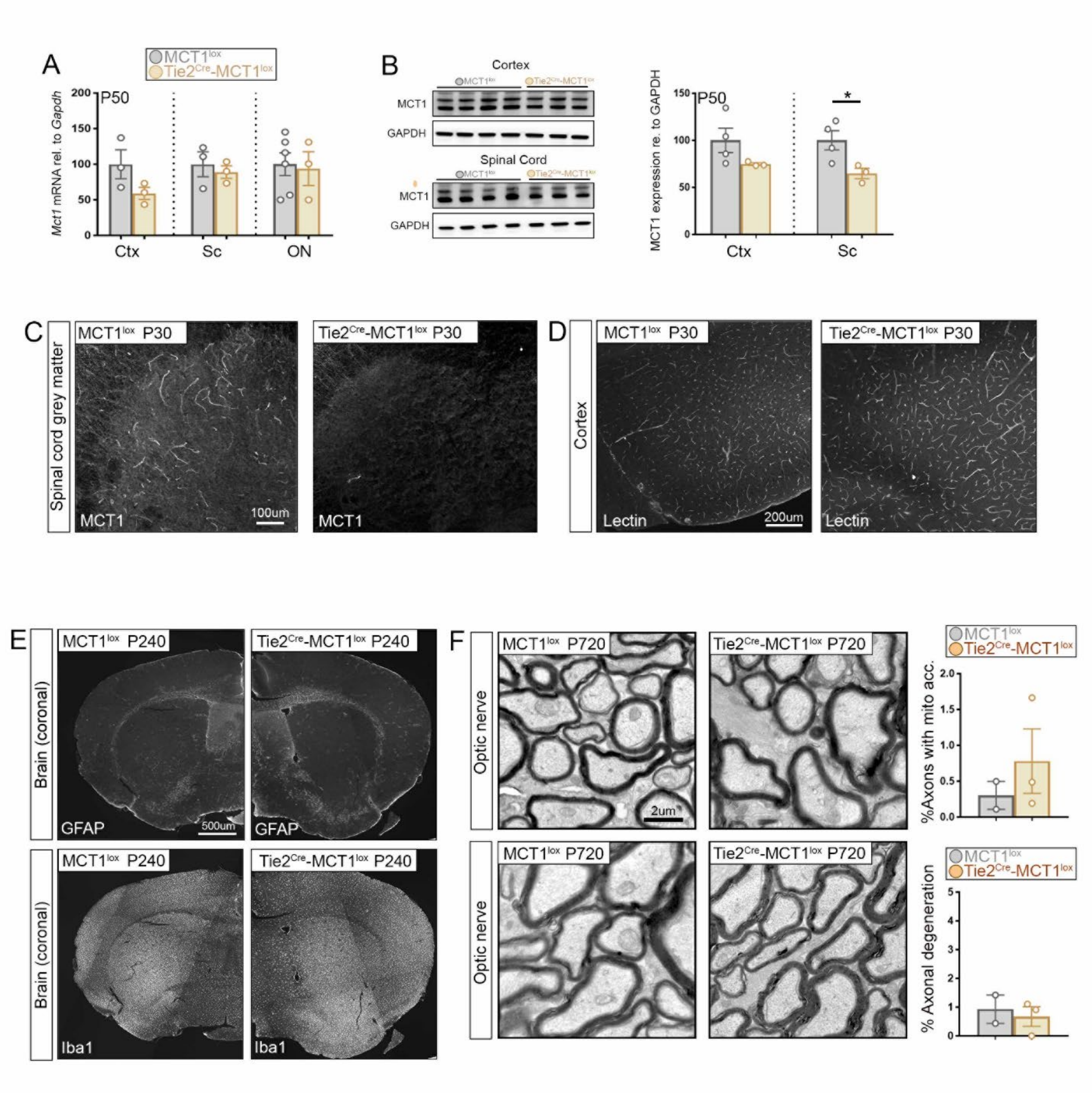

Supp Figure 5

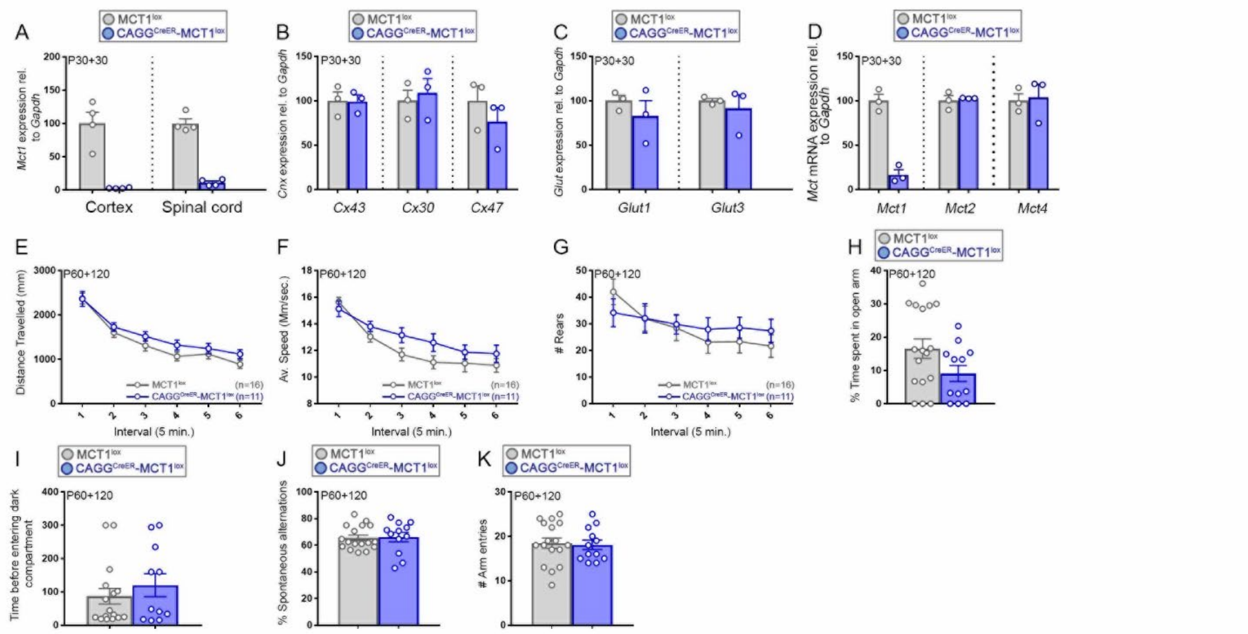

Supp Figure 6

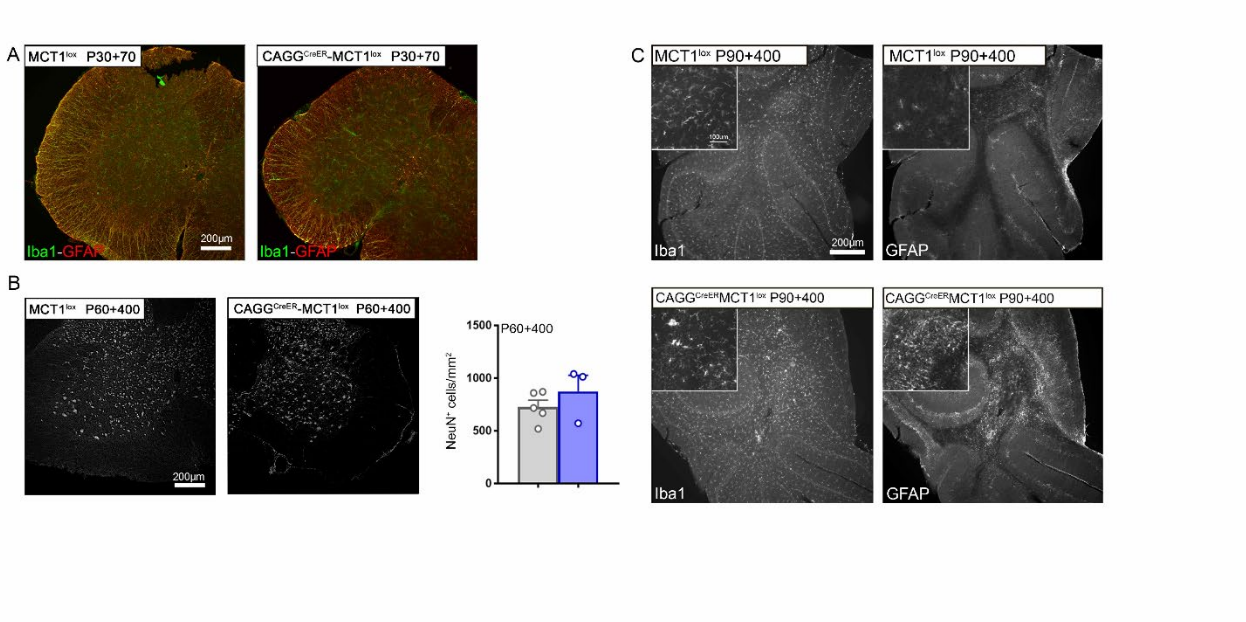
